# Impact of COVID-19 on Hospital Admission of Acute Stroke patients in Bangladesh

**DOI:** 10.1101/2020.09.28.316448

**Authors:** A T M Hasibul Hasan, Subir Chandra Das, Muhammad Sougatul Islam, Mohaimen Mansur, Md Shajedur Rahman Shawon, Rashedul Hassan, Mohammad Shah Jahirul Hoque Chowdhury, Md Badrul Alam Mondal, Quazi Deen Mohammad

## Abstract

**Background:** With the proposed pathophysiologic mechanism of neurologic injury by SARS COV-2 the frequency of stroke and henceforth the related hospital admissions were expected to rise. In this paper we investigate this presumption by comparing the frequency of admissions of stroke cases in Bangladesh before and during the pandemic.

**Methods:** We conducted a retrospective analysis of stroke admissions in a 100-bed stroke unit at the National Institute of Neurosciences and Hospital (NINS&H) which is considerably a large stroke unit. We considered all the admitted cases from the 1^st^ January to the 30^th^ June, 2020. We used Poisson regressions to determine whether statistically significant changes in admission counts can be found before and after 25 March since when there is a surge in COVID-19 infections.

**Results:** A total of 1394 stroke patients got admitted during the study period. Half of the patients were older than 60 years, whereas only 2.6% were 30 years old or younger with a male-female ratio of 1.06:1. From January to March, 2020 the mean rate of admission was 302.3 cases per month which dropped to 162.3 cases per month from April to June with an overall reduction of 46.3% in acute stroke admission per month. In those two periods, reductions in average admission per month for ischemic stroke (IST), intracerebral hemorrhage (ICH), subarachnoid hemorrhage (SAH) and venous stroke (VS) were 45.5%, 37.2%, 71.4% and 39.0%, respectively. Based on weekly data, results of Poisson regressions confirm that the average number of admissions per week dropped significantly during the last three months of the sample period. Further, in the first three months, a total of 22 cases of hyperacute stroke management were done whereas in the last three months there was an 86.4% reduction in the number of hyperacute stroke patients getting reperfusion treatment. Only 38 patients (2.7%) were later found to be RT-PCR for SARS Cov-2 positive based on nasal swab testing.

**Conclusion:** Our study revealed more than fifty percent reduction in acute stroke admission during the COVID-19 pandemic. It is still elusive whether the reduction is related to the fear of getting infected by COVID-19 from hospitalization or the overall restriction on public movement and stay-home measures.

## Introduction

Neurologic dysfunction is reported in up to one-third of the cases of COVID-19^1^. The frequency of stroke has been reported to range from 2.8% to 5.4% among confirmed and hospitalized COVID-19 patients^1,2^. Reported COVID-19-related hemorrhagic strokes are far less common than ischemic strokes^3-6^. While the pathogenesis of COVID-19-related hemorrhagic strokes is still elusive, hypercoagulable state, vasculitis and cardiomyopathy had been proposed as potential pathogenic mechanisms for ischemic stroke in COVID-19 patients^7,8^. Some researchers had stated that the viral affinity to ACE-2 receptor present in endothelium might be responsible for the rupture of intracranial vessel wall^9^. COVID-19-related stroke patients were more likely to be older, hypertensive, and had a higher D-dimer level^10^. Qin C et all^11^ reported that COVID-19 related stroke patients had more comorbidity, lower platelet counts and leukocyte counts, and the patients had higher levels of D-dimers, cardiac troponin I, NT pro-brain natriuretic peptide, and interleukin-6.

Considering the pandemic nature of the spread of COVID-19 together with the proposed pathogenic mechanism of stroke in its patient, the acute stroke cases were expected to have a sharp rise at hospitals. But in recent months neurologists from different parts of the world have reported a reasonable drop in acute stroke cases showing up at emergency care^12^. Although COVID-19 was first reported in Bangladesh on 8 March 2020 by the Institute of Epidemiology, Disease Control and Research (IEDCR), there has been a rise in the infection rate since early April. As of 15 June, the attack rate (AR) in Bangladesh is 532.1 per million^13^. For the last several weeks the case detection rate has been more than 20% and the total number of cases has exceeded the two hundred and seventy-five thousand mark on 16 August 2020^14^. As a consequence, a large number of stroke cases were also expected to get admitted to hospitals since the onset of the infection.

In this study, we assess whether the volume of actual admissions during the pandemic match suspected rise in stroke cases by conducting a trend analysis of stroke admission at the 100-bed stroke unit of the National Institute of Neuroscience & Hospital (NINS&H) which is a center of excellence, and the only center to provide comprehensive stroke care in Bangladesh.

## Materials and Methods

This was a retrospective analysis of all the admitted stroke patients at the stroke unit of NINS&H from the 1^st^ January to the 30^th^ June 2020. All the stroke cases were confirmed by CT scans and/or MRI imaging studies. Male and female patients of age 18 years or above who have confirmed diagnosis of stroke were admitted in the stroke unit. The hospital had not been declared as a Government designated COVID-19 hospital during the study period.

### The Stroke Unit and Stroke Assessment

All patients at the emergency were checked for any symptom of COVID-19 including the evaluation of chest X-ray before admission. If any patient of stroke developed shortness of breath or showed an oxygen saturation level below 90%, RT-PCR for SARS COV-2 from nasal swab were done. Those with a positive RT-PCR report were referred to the Government designated COVID-19 hospitals.

### Assessment of COVID-19 status

All patients at the emergency are checked for any feature of COVID-19 including the evaluation of chest X-ray before admission. If any patient of stroke developed shortness of breath or the oxygen saturation was less than 90%, RT-PCR for SARS CoV-2 from nasal swab were done. Those with positive RT-PCR report were referred to the Government designated COVID-19 hospitals.

### Ethical Issues

Prior to the commencement of the study, the protocol was reviewed and approved by the Ethical Review Committee (ERC), National Institute of Neurosciences and Hospital. All the data were collected manually from the hospital records. The authority provided fully anonymized data and the need for the informed consent was also waived by the ERC due to the retrospective nature of the study. This study did not include any intervention or did not impose any harm to anybody. The issues of privacy and confidentiality of patient information were strictly maintained throughout the study.

### Data Analysis

Both monthly and weekly admission rates were considered for data analysis. In case of week-wise distribution of data 12 weeks before the 25^th^ March were considered as the pre-COVID period and the 12 weeks onwards were considered as the COVID period. The chosen cut-off date of 25 March is due to the fact that, Bangladesh began to experience a substantial rise in confirmed COVID-19 cases which jumped to near 5000 from mere 39 in four weeks^13^. Data analysis was done by Statistical Package for Social Sciences (SPSS) version 21 and R version 4.0.0. In addition to showing monthly frequency distributions and time series plots, we used Poisson regression to evaluate the data for significant changes in weekly admission rates before and during the COVID-19 period.

## Results

A total of 1394 stroke patients got admitted in the stroke unit of National Institute of Neurosciences and Hospital during the study period. Among them 38 patients (2.7%) were later found to be RT-PCR for SARS Cov-2 positive based on nasal swab testing. The percentage distribution in Table 1 shows that half of the patients were older than 60 years, whereas only 2.6% were 30 years old or younger with a male to female ratio of 1.06:1.

**Table 1:** Distribution of patients by age and sex (n=1394).

| Characteristics | Groups | Number (n) | Percentage (%) |
| --- | --- | --- | --- |
| Age (in years) |  |  |  |
|  | <30 | 37 | 2.65 |
|  | 30-60 | 660 | 47.34 |
|  | >60 | 697 | 50.00 |
| Sex |  |  |  |
|  | Male | 720 | 51.60 |
|  | Female | 674 | 48.40 |

Figure 1 plots monthly time series of admissions for different types of stroke cases. A common pattern is that there is a large drop after March when the number of confirmed COVID-19 cases began to surge. From January to March (pre-COVID period) a total of 907 acute stroke cases got admitted in the stroke unit with a mean rate of admission of 302 cases per month, whereas from April to June (COVID period) the number was 487 with a mean rate of admission of 162.3 cases per month. This means an overall reduction of 46.3% in acute stroke admission per month. For all types of stroke, the number of admissions during the pre-COVID period is higher than that during the COVID period, as revealed by Figure 2. The mean admission rate for ischemic stroke, intracerebral hemorrhage, subarachnoid hemorrhage and venous stroke before and during the pandemic were 78.3 versus 42.7; 149.7 versus 94; 60.7 versus 17.3 and 13.7 versus 8.3 cases per month respectively. The reductions in monthly mean admissions for ischemic stroke (IST), intracerebral hemorrhage (ICH), subarachnoid hemorrhage (SAH) and venous stroke (VS) were 45.5%, 37.2%, 71.4% and 39.0% respectively.

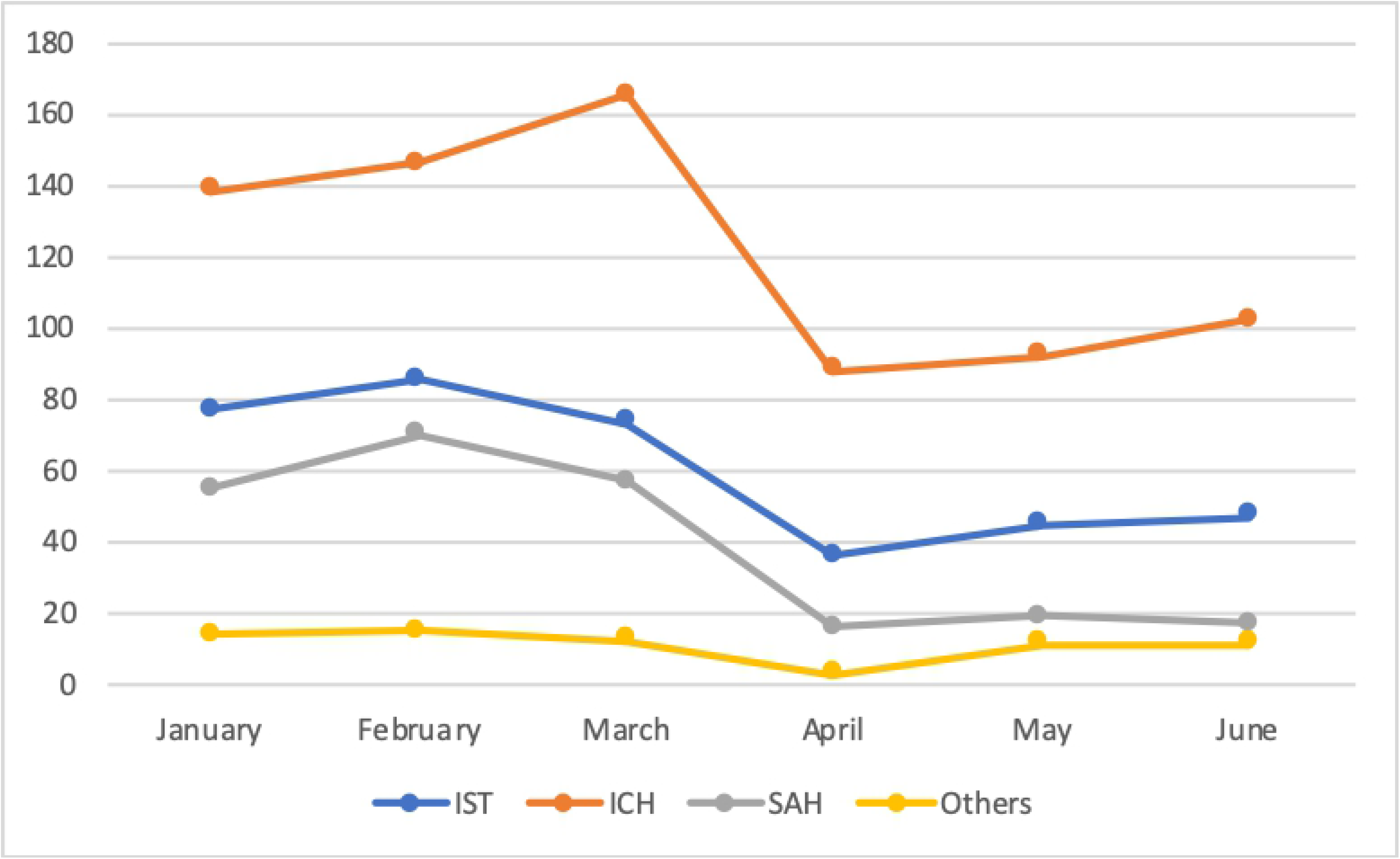

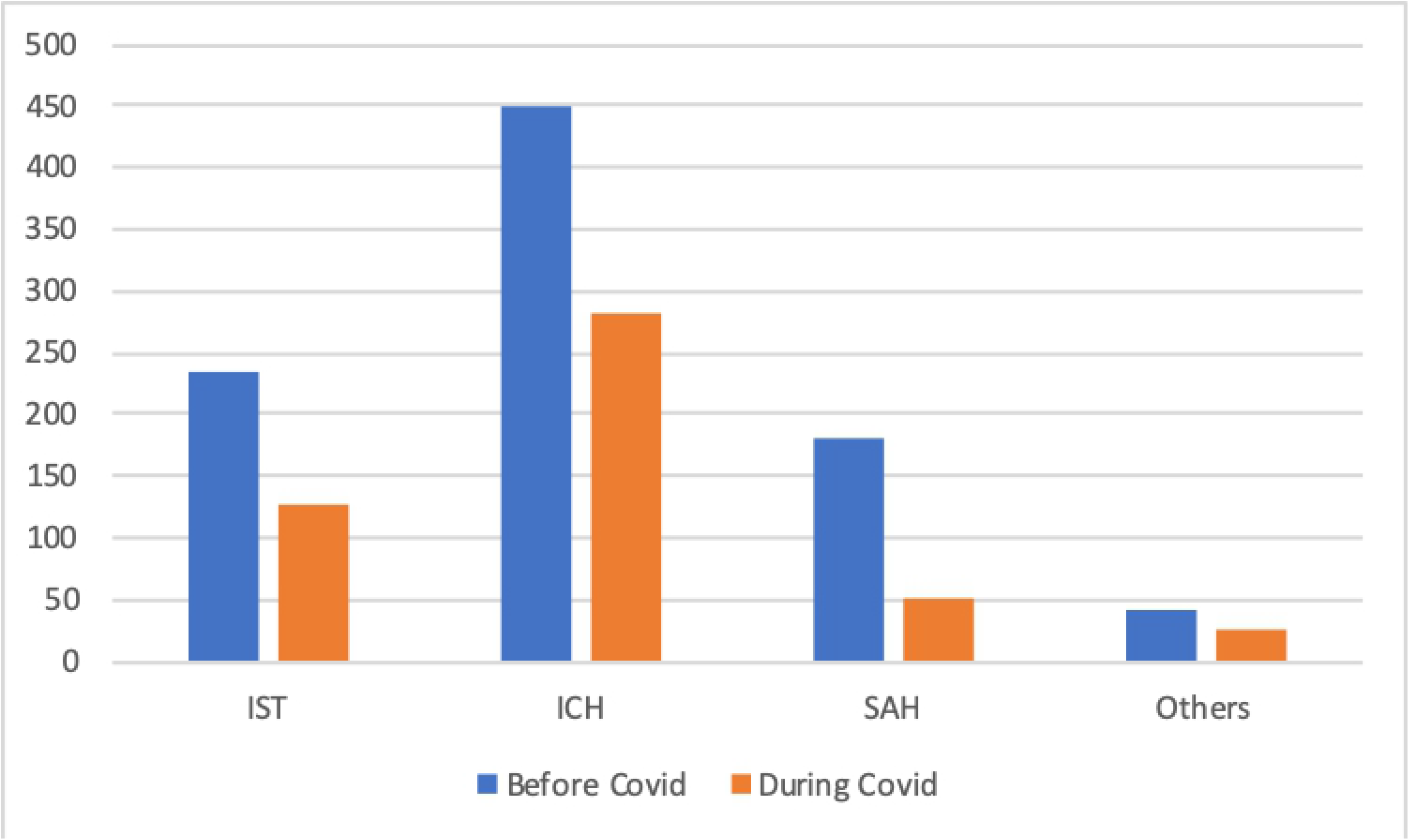

In order to get a deeper insight, we further analyze weekly admission data starting from the 1^st^ January, 2020. We considered a total of 24 weeks of which the first 12 weeks is considered as the pre-COVID period and the last 12 weeks as the COVID period. For each stroke type, we run a Poisson regression where the dependent variable is the count of admissions of stroke patients and the independent variable is a COVID period indicator (0 = pre-COVID, 1 = COVID). Results of the regression are reported in Table 2. Negative coefficients with p-values < 0.05 indicate statistically significant drops in the weekly rates of all types of stroke admissions during the COVID period.

**Table 2:**
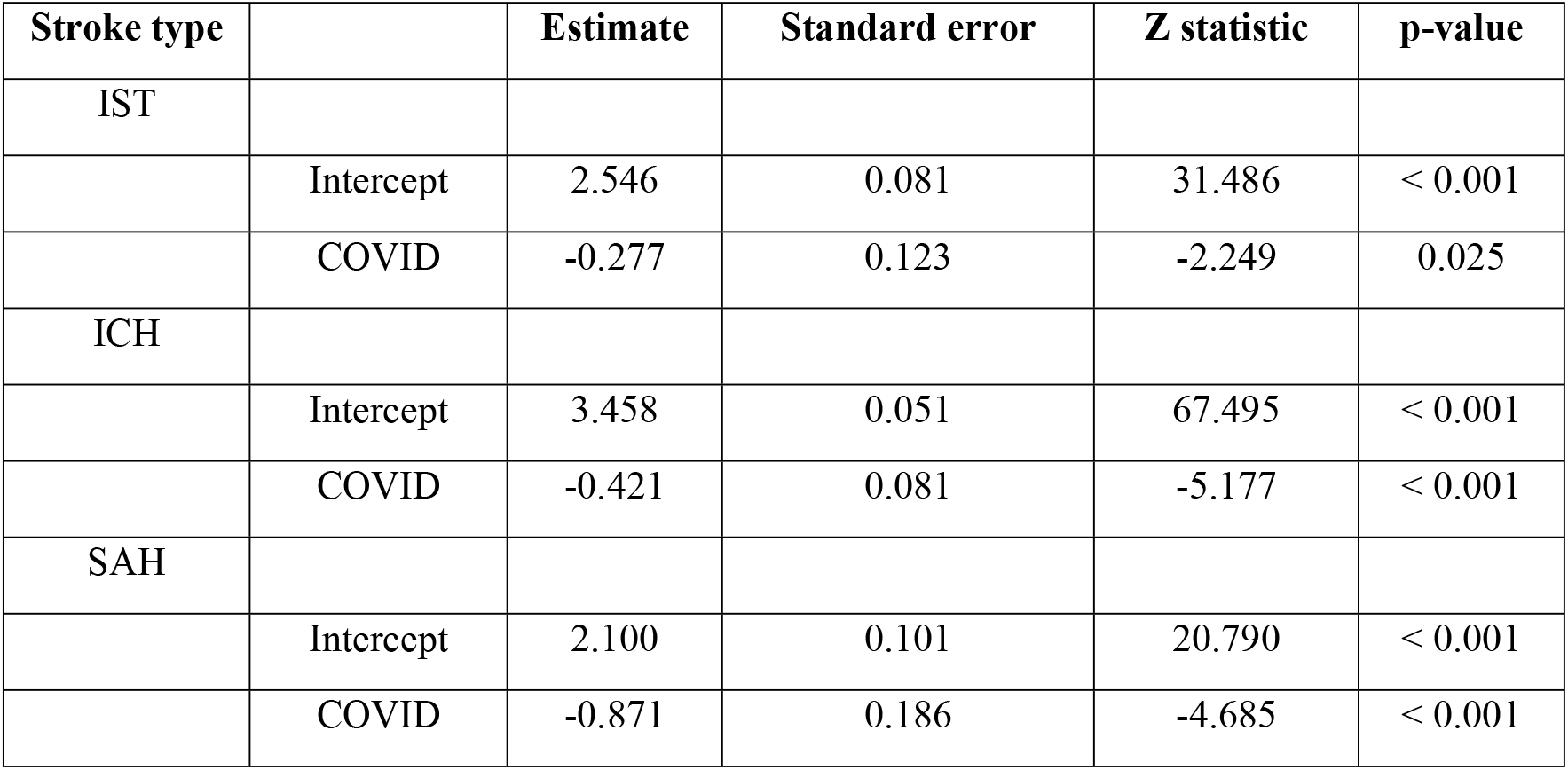
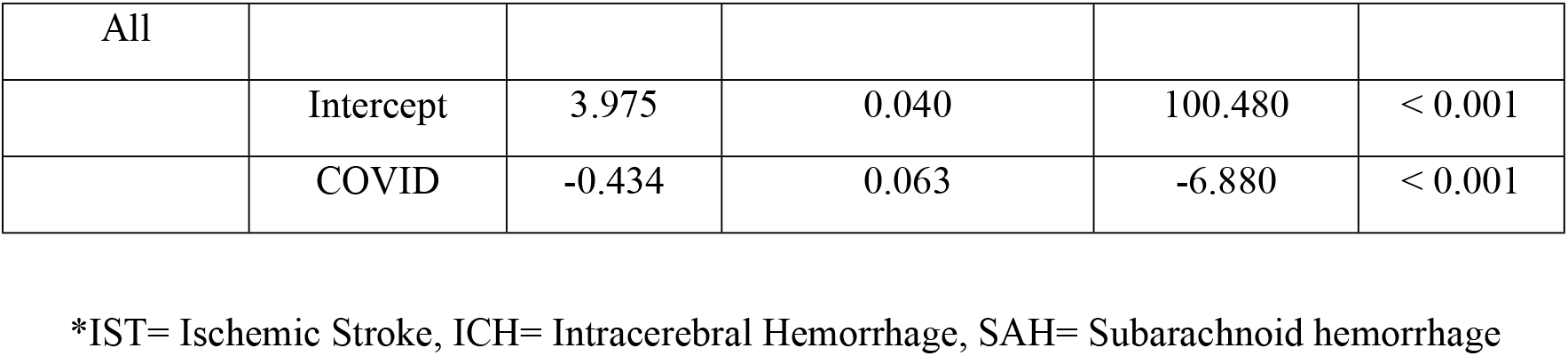
Results of Poisson regressions for different stroke types

| Stroke type |  | Estimate | Standard error | Z statistic | p-value |
| --- | --- | --- | --- | --- | --- |
| IST |  |  |  |  |  |
|  | Intercept | 2.546 | 0.081 | 31.486 | < 0.001 |
|  | COVID | -0.277 | 0.123 | -2.249 | 0.025 |
| ICH |  |  |  |  |  |
|  | Intercept | 3.458 | 0.051 | 67.495 | < 0.001 |
|  | COVID | -0.421 | 0.081 | -5.177 | < 0.001 |
| SAH |  |  |  |  |  |
|  | Intercept | 2.100 | 0.101 | 20.790 | < 0.001 |
|  | COVID | -0.871 | 0.186 | -4.685 | < 0.001 |
| All |  |  |  |  |  |
|  | Intercept | 3.975 | 0.040 | 100.480 | < 0.001 |
|  | COVID | -0.434 | 0.063 | -6.880 | < 0.001 |

\*IST= Ischemic Stroke, ICH= Intracerebral Hemorrhage, SAH= Subarachnoid hemorrhage

We check robustness of our regression results by further conducting a statistical test of difference between two Poisson rates modified for small sample applications^15^. The null hypothesis of the test is that there is no difference in the weekly rate of stroke admissions during the pre-COVID period, λ_1_ and the weekly rate during the COVIID period, λ_2_, and the alternative hypothesis is that λ_2_ < λ_1_. Results of the tests reported in Table 3 confirm rejection of the null hypothesis at least at 5% level of significance and support the finding of the Poisson regression that the weekly rate of admission for all types of stroke patients reduced significantly during the COVID period compared to the pre-COVID period.

**Table 3:** Results of a test for difference in the rates of weekly admission before and during the COVID period

| Stroke type | A test for the difference of two Poisson rates |  |
| --- | --- | --- |
| | $H_0: \lambda_1 = \lambda_2$ vs $H_1: \lambda_2 < \lambda_1$ | |
|  | Z statistic | p-value |
| IST | 2.26 | 0.012 |
| ICH | 5.24 | < 0.001 |
| SAH | 4.94 | < 0.001 |
| All | 6.97 | < 0.001 |

\*IST= Ischemic Stroke, ICH= Intracerebral Hemorrhage, SAH= Subarachnoid hemorrhage

In the first three months a total of 21 cases of hyperacute stoke management was done in the form of I/V thrombolysis in the stroke unit while in the last three months we observed an 85.7% reduction in the number of hyper acute stroke patients receiving I/V thrombolysis and reperfusion treatment. This downward trend is captured in Figure 3.

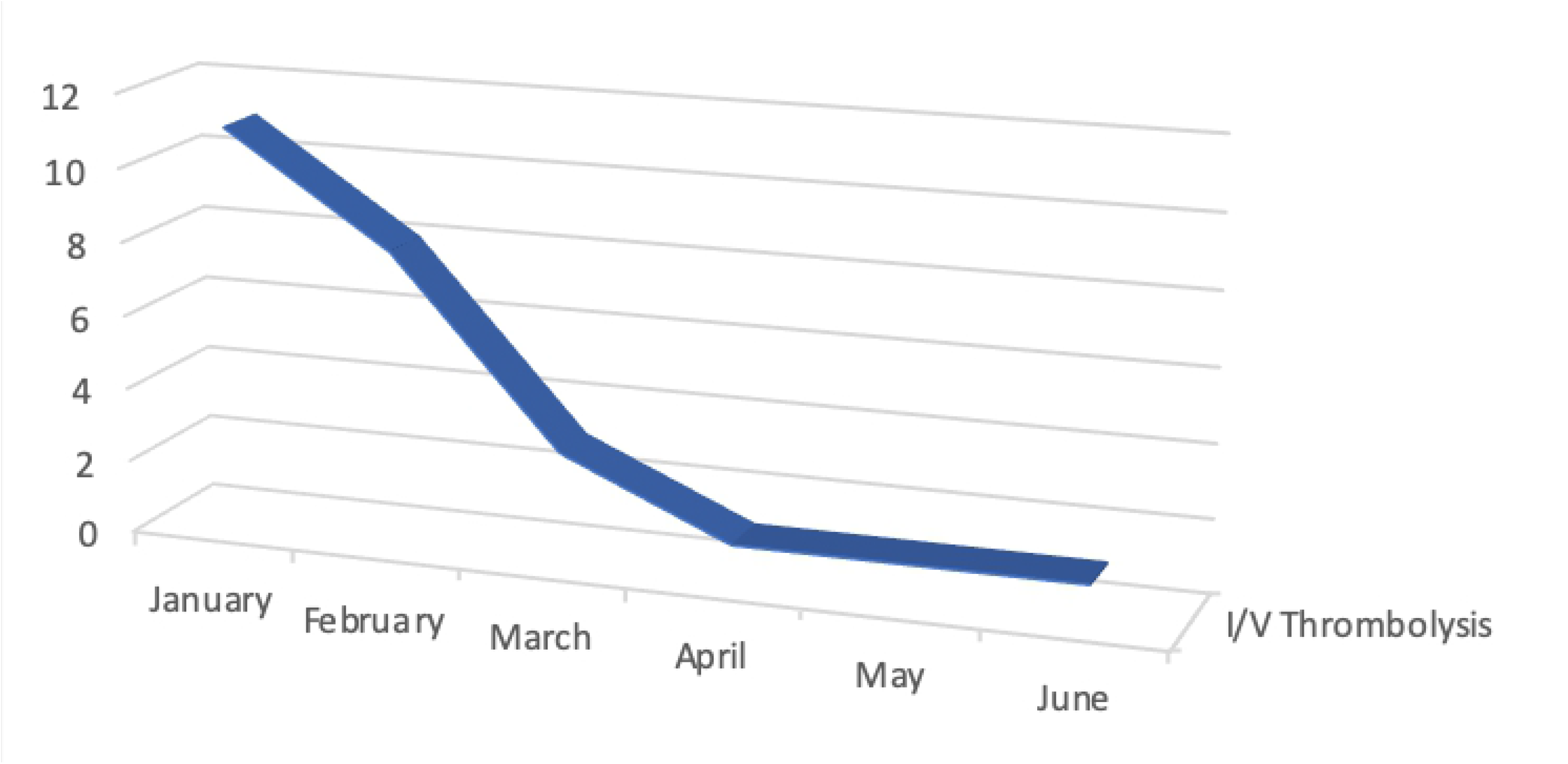

## Discussion

Our study provides an insight into the impact of COVID-19 on admission of acute stroke patients in Bangladesh. As the pandemic hit the country, several hospitals were designated by the Government exclusively for the treatment of COVID-19 patients. These include the Dhaka Medical College Hospital which deals with a considerable portion of stroke cases each year. For this reason, the stroke unit of National Institute of Neurosciences and Hospital was expected to experience a higher than normal patient load during the spread of the pandemic.

With the report of several cases of large vessel occlusion from New York and taking the pathogenic mechanism of neurologic syndromes in COVID-19 patients into consideration, the number of acute stroke cases in Bangladesh was expected to show a steep rise during the pandemic^2,7-10^. In contrary to the forecast, our analysis based on number of admitted stroke cases in the largest stroke treatment center of the country over the period of January to June, 2020 revealed a gross (46.3%) reduction in the rate of acute stroke admissions in last three months of the study period when the infection surged. The reduction for SAH admissions was the highest (71.4%) followed by IST (45.6%), VS (39.02%) and ICH (37.1%) admissions. Results of Poisson regressions confirm that reductions in weekly admission rates during the COVID period are statistically significant. Hoyer et al^16^ reported a similar picture of acute stroke admission during the pandemic based on data from four academic stroke centers in Germany. They compared stroke admission rates at the same weeks of the year 2019 and 2020, and reported 38- 46% drop in acute stroke admissions in those centers. They assumed that while factors such as size of stroke center’s catchment area may be responsible for differential reductions in the four centers, the fear of getting infected at hospitals and forced lock-down measures may have resulted in lower interest in seeking medical help and consequent overall reduction in stroke admissions^16^. With agreement to Dhand A et al^17^ he also opined that social distancing and confinement at home might have resulted in delayed symptoms disclosure and late or no hospital admission with wait-and-watch strategy.

Moreover, COVID-19 related stroke admissions in our study were only 2.7% which was similar to the report of Qin C et al^11^ from Wuhan, China but three times of that reported by Yaghi S et al^18^ from New York. Bangladesh did not impose any nation-wide lock down following the detection of its first COVID-19 case on the 8^th^ of March, 2020. Only on the 26^th^ of March a nation-wide public holiday was declared and the public transport was placed under restriction. These measures were subsequently extended for several weeks. However, movements of ambulance or other emergency services were exempted from these restrictions. Therefore, it is unlikely that the reduction in admitted stroke cases is due to the restriction on movements or closure of all type of public transports. It is also not clear whether the fear of getting infected at hospitals or the overall ‘stay home’ publicity in general is the main cause of this reduction.

We had a few limitations in this study. First of all, this is a retrospective analysis based on hospital records. Therefore, available information were of limited value and further analyses such as investigating reasons behind the reduction in stroke admissions were not possible. Secondly, since the stroke unit at NINS&H started its journey only in September 2019, we could not compare stroke admissions during 2020 with the same time periods of the year 2019, which would have been a more fare comparison devoid of any possible seasonal effects on admission. The major strength of the study, however, is that we could analyze a considerable number of admitted stroke cases over substantially long periods before and during the COVID-19 pandemic. Importantly, the data came from one of the largest stroke units in the world and the highest center of referral for stroke cases in Bangladesh that provides a comprehensive stroke care 24×7.

## Conclusion

In contrary to the presumption of expecting a higher stroke burden in hospitals during the COVID-19 pandemic our study found more than fifty percent reduction in the number of acute stroke admissions in the largest stroke unit in Bangladesh. The decline was the most prominent in SAH admissions followed by IST admissions. Thus, this study is another addition to the recent evidence of such decreasing trends in stroke admissions during the pandemic. Whether the reduction is related to the fear of getting infected by COVID-19 from hospitalization or the overall restriction of public movement and stay-home measures needs to be investigated by further studies. Even in this time of the pandemic and consequent social distancing measures, it is crucial for patients experiencing acute stroke attacks to seek immediate medical help so that timely diagnosis, intervention and management of stroke is possible. Increasing public awareness in this regard will help avoid unnecessary and unfortunate losses of lives from untreated strokes.

## Authors Contribution

Dr. ATM HH was involved in planning the study, setting the methodology, consultation and data collection and write up for this study. Dr. SCD was involved in data collection and patient management. Dr. MSI, Dr. MM and Dr. MSRS were involved in data analysis, data interpretation and partly writing the manuscript. The rest were involved in consultation and data collection. All the authors have read and approved the final version of the manuscript.

## Acknowledgement

We acknowledge the contribution of our supporting staff at the stroke unit, NINS&H for helping us with the necessary data for the study.

